# Widespread RNA editing dysregulation in Autism Spectrum Disorders

**DOI:** 10.1101/446625

**Authors:** Stephen Tran, Hyun-Ik Jun, Jae Hoon Bahn, Adel Azghadi, Gokul Ramaswami, Eric L. Van Nostrand, Thai B. Nguyen, Yun-Hua E. Hsiao, Changhoon Lee, Gabriel A. Pratt, Gene W. Yeo, Daniel H. Geschwind, Xinshu Xiao

## Abstract

Autism spectrum disorder (ASD) is a genetically complex, clinically heterogeneous neurodevelopmental disease. Recently, our understanding of the molecular abnormalities in ASD has been expanded through transcriptomic analyses of postmortem brains. However, a crucial molecular pathway involved in synaptic development, RNA editing, has not yet been studied on a genome-wide scale. Here, we profiled the global patterns of adenosine-to-inosine (A-to-I) editing in a large cohort of post-mortem ASD brains. Strikingly, we observed a global bias of hypo-editing in ASD brains, common to different brain regions and involving many genes with known neurobiological functions. Through genome-wide protein-RNA binding analyses and detailed molecular assays, we show that the Fragile X proteins, FMRP and FXR1P, interact with ADAR proteins and modulate A-to-I editing. Furthermore, we observed convergent patterns of RNA editing alterations in ASD and Fragile X syndrome, thus establishing RNA editing as a molecular link underlying these two highly related diseases. Our findings support a role for RNA editing dysregulation in ASD and highlight novel mechanisms for RNA editing regulation.

## Introduction

Autism spectrum disorder (ASD) is a neurodevelopmental deficit in social communication accompanied by repetitive and restrictive interests^1^. ASD pathology implicates many aspects of nervous system biology including dysregulation in glutamatergic^2,3^ and serotonergic^4-6^ circuits, aberrant structural development in multiple brain regions^7^, excitatory and inhibitory imbalance^8^, and abnormal synaptogenesis^9,10^. However, the genetic etiology of ASD remains incompletely understood and shows substantial heterogeneity^6,11,12^. Nevertheless, recent studies, taking advantage of the increasing availability of postmortem samples, have revealed shared patterns of transcriptome dysregulation across approximately 2/3 of ASD patients affecting neuronal and glial coding and non-coding gene expression, neuronal splicing^13-15^ including microexons^16,17^, and microRNA targeting^18^. These studies highlight biological pathways, involving down-regulation of activity-dependent genes in neurons and up-regulation of astrocyte and microglial genes, as key points of convergence in ASD pathology.

Another major RNA processing mechanism is RNA editing. RNA editing refers to the alteration of RNA sequences through insertion, deletion or substitution of nucleotides^19^. Catalyzed by the ADAR family of enzymes, adenosine-to-inosine (A-to-I) editing is the most prevalent type of RNA editing in humans, affecting the majority of human genes^19,20^. As inosines in the RNA are recognized as guanosines by cellular machinery, A-to-I editing can alter gene expression in different ways, for example, through amino acid substitutions, modulation of RNA stability, alteration of alternative splicing, and modifications of regulatory RNAs or *cis*-regulatory motifs^21,22^.

RNA editing plays important roles in neurodevelopment and maintenance of normal neuronal function^21,23^. A number of A-to-I editing sites have been identified as classical examples of RNA editing in modulating excitatory responses and permeability of ionic channels and other neuronal signaling functions^21^. Not surprisingly, aberrant RNA editing has been reported in several neurological disorders, such as schizophrenia^24^, bipolar disorder^25^, Alzheimer’s disease^26,27^ and amyotrophic lateral sclerosis^28,29^. In ASD, a previous study analyzed a few known RNA editing sites in synaptic genes and reported altered editing patterns in a small cohort of ASD cerebella^30^. Yet, it remains to be understood if global patterns of RNA editing may contribute to the neuropathology of ASD, a question that calls for expanded studies of large patient cohorts and multiple implicated brain regions. In addition, the regulatory mechanisms of aberrant editing in neurological disorders including ASD remain largely unknown.

Here we report global patterns of dysregulated RNA editing across the largest cohort of ASD brain samples to date, spanning multiple brain regions. We identified a core set of down-regulated RNA editing sites, enriched in genes of glutamatergic and synaptic pathways and ASD susceptibility genes. Multiple lines of evidence associate a distinct set of these hypoedited sites with Fragile X proteins: FMRP and FXR1P. Through genome-wide protein-RNA binding analyses and detailed molecular assays, we showed that FMRP and FXR1P interact with ADAR and modulate A-to-I editing. It is known that mutations in FMRP lead to the Fragile X syndrome, a disease with high comorbidity with ASD^31^. Indeed, we observed convergent dysregulated patterns of RNA editing in Fragile X and ASD patients, which is consistent with the findings that genes harboring ASD risk mutations are enriched in FMRP targets^32,33^. Overall, we provide global insights regarding RNA editing in ASD pathogenesis and elucidate a regulatory function of Fragile X proteins in RNA editing that additionally serves as a molecular link between ASD and the Fragile X Syndrome.

## Results

### RNA editing analysis of ASD postmortem brain samples

We obtained a total of 261 RNA-Seq datasets derived from samples representing three brain regions implicated in ASD-susceptibility: frontal cortex, temporal cortex, and cerebellar vermis (Table S1). These datasets were generated as part of our transcriptomic study of ASD brains that included a cohort of 68 ASD and control subjects^15^. Overall, the ASD and control groups did not have significant difference in variables that could confound RNA editing analysis, such as age, gender, sequencing depth, etc (Fig. S1). For each brain sample, rRNA-depleted total RNA was sequenced with an average of 70 million raw read pairs (average 55 million uniquely mapped pairs, 2x50 nt, non-strand-specific, Fig. S2)^15^.

We applied our previously developed methods to identify RNA editing sites using the RNA-Seq data^34,35^. In addition, we implemented additional steps to capture editing sites located in “hyperedited” regions, which were likely missed by regular methods^36^ (Methods). Combining these approaches, we identified a total of 98,477 RNA-DNA differences (RDDs) from frontal cortex, 97,994 from temporal cortex, and 134,085 from cerebellum. As expected, the number of RDDs per sample correlated with read coverage approximately (Fig. S3).

On average, >95% RDDs were A-to-G and T-to-C editing types per sample, consistent with A-to-I editing reflected in non-strand-specific RNA-Seq data (Fig. 1a, Fig. S4a). The remaining 5% of RDDs mainly consisted of C-to-T and G-to-A types, possibly due to C-to-U editing. Notably, most (84%) of the A-to-I editing sites identified here are included in the REDIportal database^37^ (Fig. S4b). As expected, the majority of RDDs were located in Alu sequences^37^ (Fig. S4c) and in intronic regions^37^ (Fig. S4d). The sequence context of A-to-G sites was consistent with the typical sequence signature known for ADAR substrates^38-40^ (Fig. S4e). Together, these results strongly supported the validity of our predicted A-to-I editing sites.

**Figure 1.**
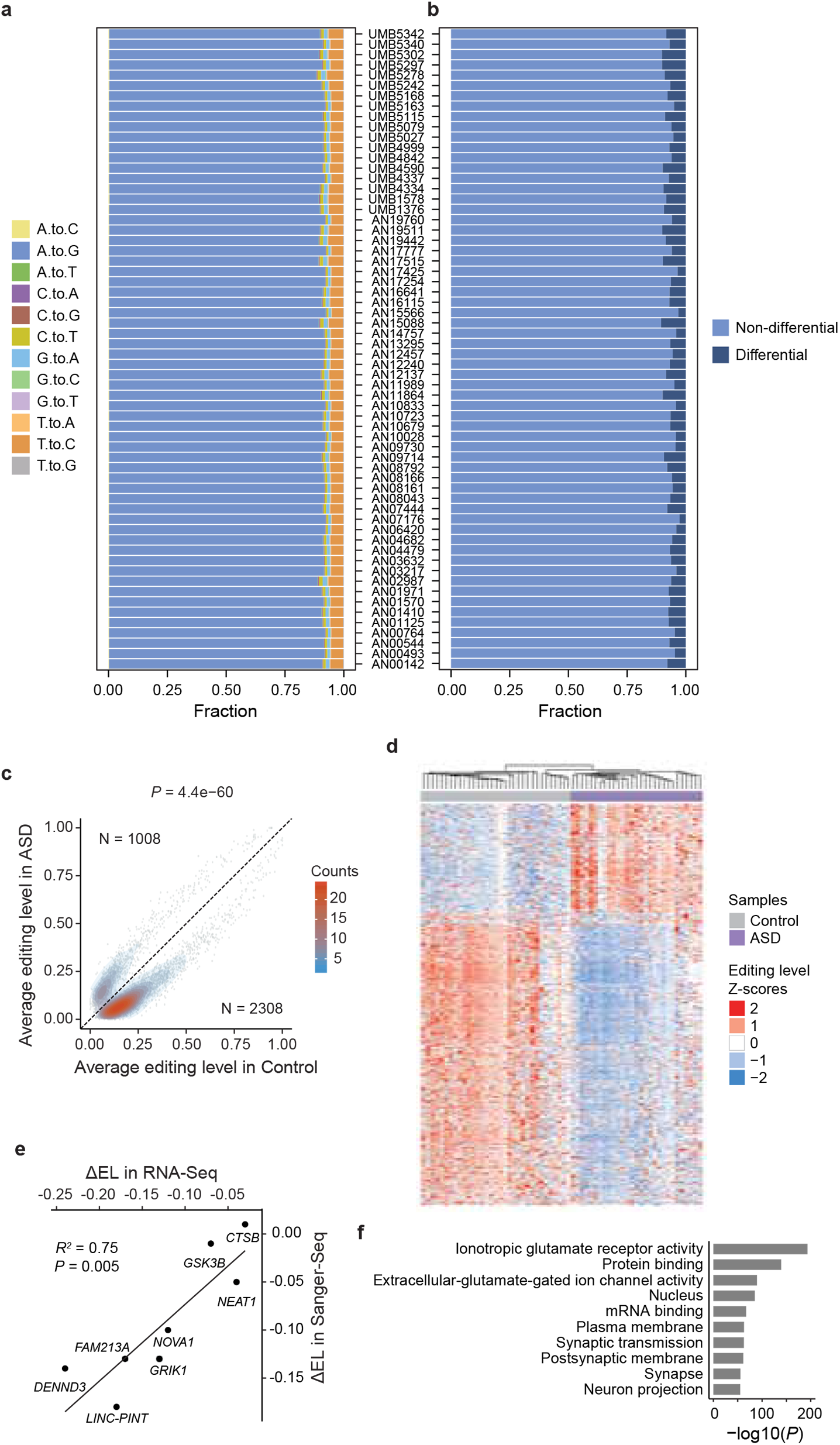
Transcriptome-wide differential editing in the frontal cortex of ASD. a, Fraction of all types of RNA-DNA differences (RDDs) identified in the RNA-Seq data of each subject. b, Fraction of differential and non-differential editing sites for each subject. c, Average editing levels of differential editing sites in ASD and controls. Numbers of editing sites that were up- and down-regulated in ASD are shown, which were compared via Fisher’s Exact test (P value shown above plot). d, Differential editing sites segregate ASD and control samples. Normalized editing levels (z-scores) were used in hierarchical clustering. Each row corresponds to one editing site. Each column represents a sample. e, Experimental validation of differential editing levels using Sanger sequencing. The frontal cortex samples used in this experiment are shown in Table S1. ΔEL: change in editing level (AΔSD - control). f, GO enrichment analysis of genes harboring differential editing sites.

Among all predicted RNA editing sites, the frontal and temporal cortex shared more than 70% of their sites, while the two cortical regions and cerebellum shared 50-55% (Fig. S4f). Furthermore, the editing levels of common editing sites between two brain regions were highly consistent (correlation coefficient 0.96 between cortex, and 0.89 to 0.90 between cortex and cerebellum, Fig. S4g). Thus, the three brain regions demonstrated similarities and differences in RNA editomes. Notably, the lower correlation between cortex and cerebellum likely reflects the substantial differences in cellular composition and physiology between these two regions^41^.

### Reduction of RNA editing in ASD frontal cortex

Given the observed difference in RNA editing between brain regions, we first focused on analysis of RNA editing dysregulation in frontal cortex, a region previously shown to have strong transcriptomic alterations in ASD^13,15^. Comparing the editing profiles of ASD and controls, we identified a total of 3,314 differential editing sites (p < 0.05, and editing level difference ≥ 5% or editing prevalence difference ≥ 10%, see Methods and Table S2). For each individual, 2.6-10.5% of all editing sites were identified as differential sites (Fig. 1b). Importantly, the differentially edited sites showed a striking bias of hypoediting in ASD samples (Fig. 1c). The number of down-regulated RNA editing sites in ASD far outnumbered those that were upregulated (p = 4.4e-60, Fisher’s Exact test, Fig. 1c).

To further confirm that the observed hypoediting pattern did not reflect possible bias in certain confounding variables, we carried out a principal component analysis (PCA) of differential editing sites and correlated the PCs with a series of biological and technical variables. Remarkably, diagnosis (i.e., ASD or control) was the only variable that demonstrated a significant association with the first PC (Fig. S5a). Consistent with this result, differential editing sites largely segregated the two groups of subjects (Fig. 1d). In addition, genes with differential editing sites had very small average gene expression difference between ASD and control groups (Fig. S5b). Thus, differential editing observed here was unlikely secondary to differential gene expression.

We carried out Sanger sequencing to confirm the observed editing difference of 8 editing sites (Table S3). These editing sites were chosen to cover a range of editing level differences (Fig. 1e). Each editing site was tested in eight postmortem frontal cortex samples (4 ASD, 4 controls) that were selected based on sample availability (Table S1b). The editing differences calculated from RNA-Seq strongly correlated with those based on Sanger sequencing (Fig. 1e, R^2^ = 0.75), confirming the accuracy of our editing level quantification.

A total of 1,189 genes were identified harboring at least one differential editing site in frontal cortex. These genes exhibited significant gene ontology (GO) enrichment for categories such as ionotropic glutamate receptor activity, glutamate gated ion channel activity, and synaptic transmission (Fig. 1f). Consistent with this finding, genes (e.g., *KCNIP4*, *PCDH9*, *RBFOX1*, and *CNTNAP2*) with the largest number of differential editing sites (either before or after correction for gene length, Fig. S6a, b) are often involved in the above functional categories. Importantly, a number of genes with differential editing were known ASD susceptibility genes^42^ (Fig. S6c). Together, these results indicate that RNA editing could contribute towards aberrant synaptic formation in ASD.

### Global analysis of potential regulators of hypoediting in ASD

To understand the regulatory mechanisms of hypoediting in ASD brains, we first examined the mRNA and protein expression levels of the *ADAR* genes. However, we did not observe significant differences of *ADAR* expression between ASD and control frontal cortex (Fig. 2a-c). In addition, the *ADAR* genes did not exhibit differential alternative splicing in these samples, as determined by our previous study^15^. With regards to genomic variation in ADAR, no studies to date have reported either rare or common variants associated with ASD in this gene.

**Figure 2.**
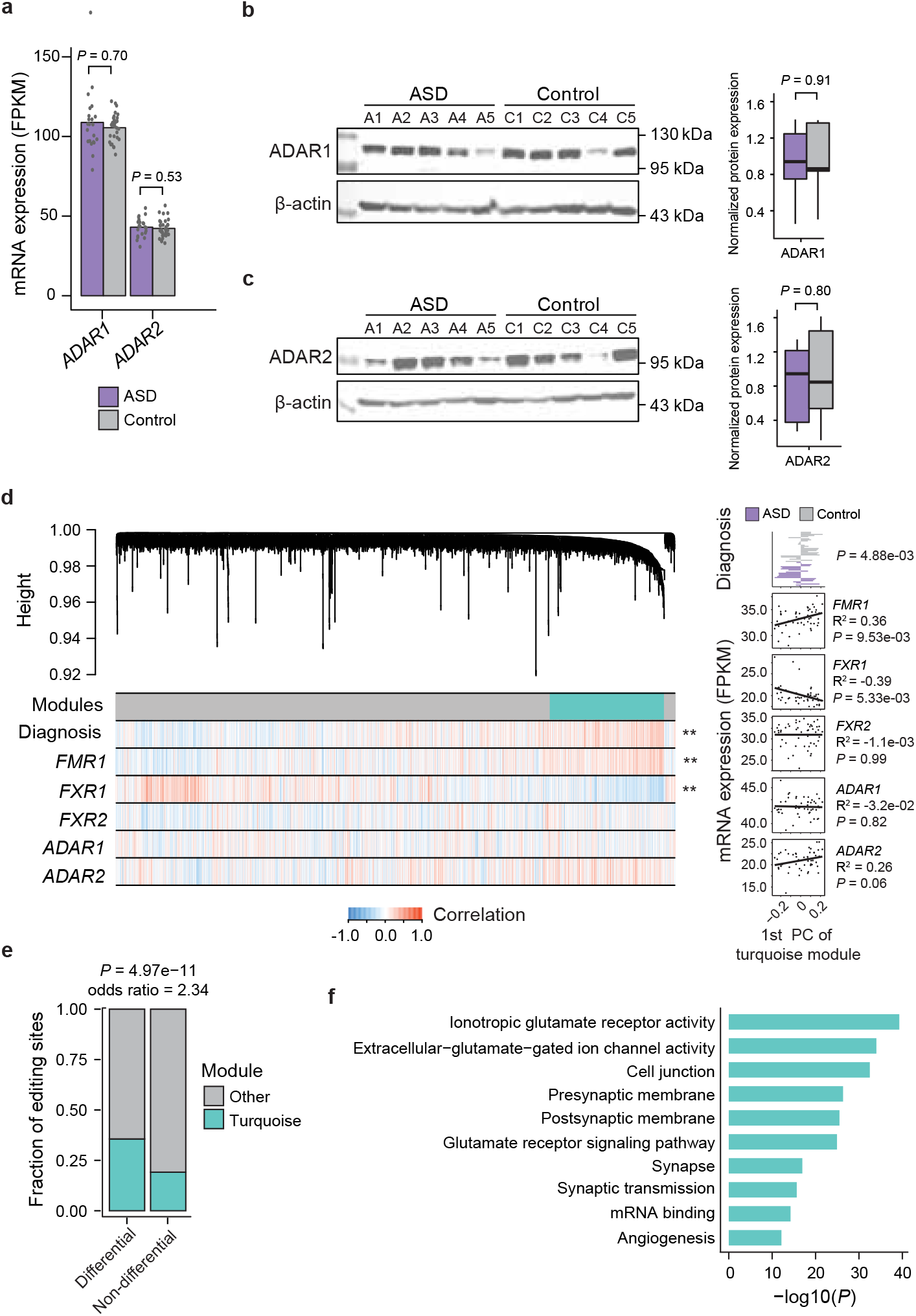
Global analysis reveals potential regulators of differential editing in the frontal cortex of ASD. a, mRNA expression levels (FPKM) of ADAR1 and ADAR2 estimated from RNA-Seq data. P values were calculated using a regression approach where covariates were accounted for (Parikshak et al, Nature, 2016). Error bars represent standard errors (same below). b, Western blot of ADAR1 protein in ASD and control samples. Protein level was normalized against that of β-actin. Samples used in this experiment are shown in Table S1 (chosen based on availability). A1-A5: ASD samples. C1-C5: control samples. P value was calculated by Student’s t-test. c, Similar as b, for ADAR2 protein. d, WGCNA analysis of RNA editing in frontal cortex. Dendrogram of RNA editing sites is shown. The turquoise module is indicated by the turquoise color. Correlation of editing sites with diagnosis (ASD or control) and mRNA expression levels of a few proteins is shown in the Heatmap. **: P < 0.01. Right panels: Bar graph and scatter plots represent association between diagnosis or mRNA expression levels and the first principal component (PC) of the turquoise module. P values of Pearson’s correlation are shown. e, Overlap between the turquoise sites and differential editing sites. P value was calculated by Fisher’s Exact test. f, GO enrichment analysis of genes harboring the turquoise sites.

We hypothesized that given the absence of explanatory variation in *ADAR* in ASD, other trans-regulators must contribute to the dysregulation of RNA editing in ASD. Given the large-scale editome profiles in this study, we reasoned that if a prevailing mechanism exists for hypoediting in ASD, then a significant number of editing sites should demonstrate correlated variation across the subjects. We applied weighted gene co-expression network analysis (WGCNA)^43^ to search for highly correlated clusters of editing sites (i.e., modules) (Methods).

Remarkably, we identified a module enriched in editing sites that had significant association with diagnosis based on the correlation of editing variation across subjects (Fig. 2d, Table S4). We next asked whether certain trans-regulators were responsible for regulation of this module (called the “turquoise module”) by examining the correlation between the module “eigengene”^44^ (i.e., eigen-editing site) and expression levels of potential trans-regulators. We included a small number of RNA-binding proteins (RBPs) in the search of trans-regulators chosen based on a parallel study (Tran, Bahn, Xiao, unpublished data). Interestingly, we identified strong association between the turquoise module and Fragile X-relevant genes (*FMR1* and *FXR1*) (Fig. 2d). *FMR1* associated with editing changes were positively correlated, reflecting possible enhancing function in editing regulation. In contrast, *FXR1* demonstrated a negative association, indicating possible repressive regulation. Importantly, the turquoise module was enriched with differential editing sites between ASD and controls (Fig. 2e), suggesting that the Fragile X-related genes may be responsible for hypoediting in ASD. In addition, this module is significantly enriched with genes related to synaptic ontology (Fig. 2f), consistent with a primary known function of FMRP in localization and maintenance of synaptic membranes^45,46^, and previous reports of FMRP binding targets being enriched in ASD risk genes^32,33^.

### Interaction between Fragile X proteins and ADARs

To further examine the involvement of Fragile X proteins in RNA editing regulation, we asked whether these proteins were co-localized or interacted with ADAR proteins. In the literature, FMRP is known to predominantly reside in the cytoplasm^47,48^, although it has also been shown to bind to RNAs in the nucleus^49^. In contrast, ADAR1 and ADAR2 mainly reside in the nucleus^22^. To confirm these findings, we carried out a subcellular fractionation experiment followed by Western blot in HeLa cells. Consistent with previous literature, the ADAR proteins were abundant in the nuclear fraction and FMRP and FXR1P were detected in the cytoplasmic fraction (Fig. 3a). Nevertheless, the Fragile X proteins were also abundant in the nucleus. The subcellular distribution of ADAR proteins remained unchanged upon FMRP or FXR1P knockdown (Fig. 3a).

**Figure 3.**
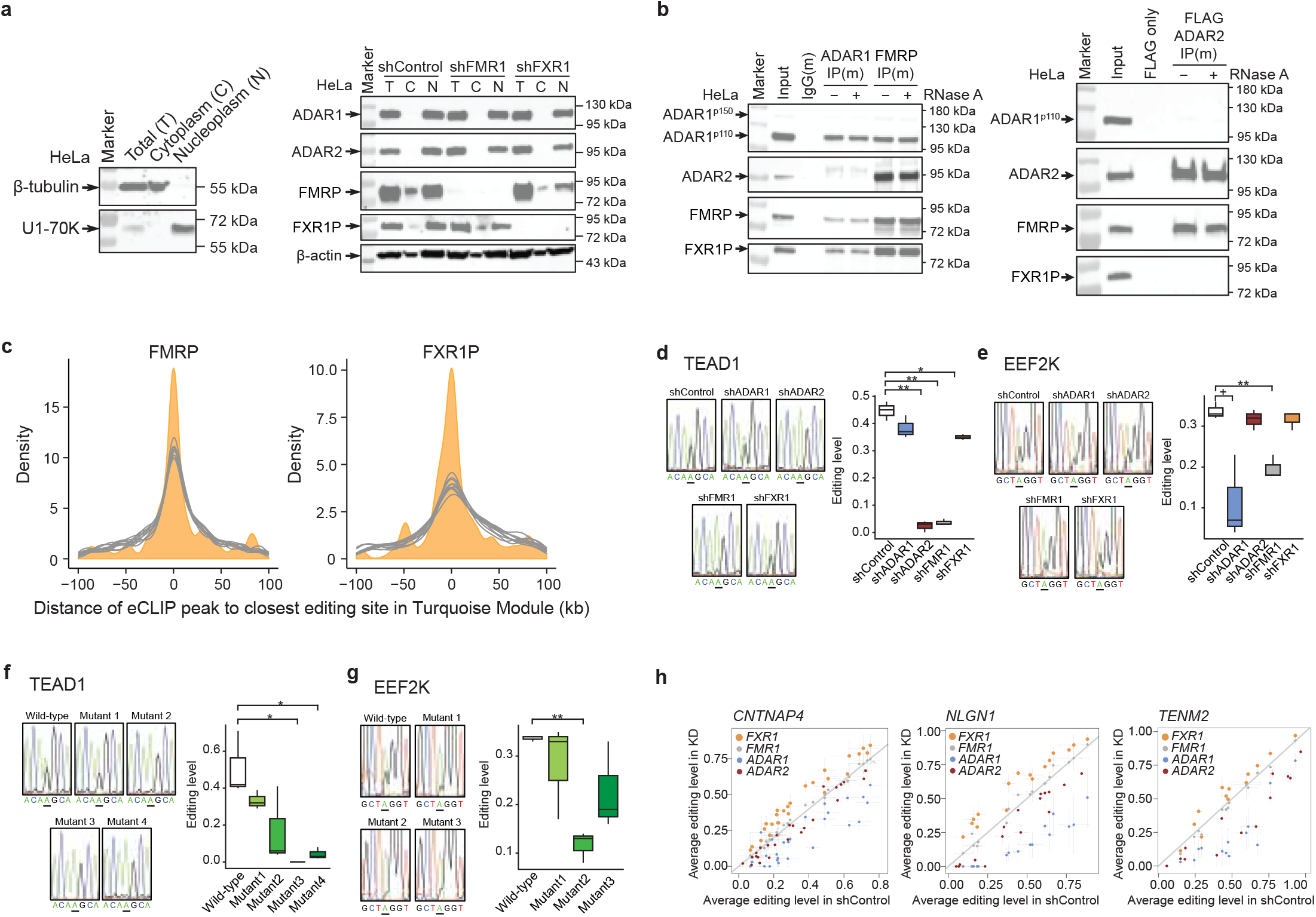
FMRP and FXR1P regulate RNA editing. a, Western blot of ADAR1, ADAR2, FMRP and FXR1P proteins in the nuclear and cytoplasmic fractions of HeLa cells. Cell fractionation was confirmed by Western blotting of β-tubulin and U1-70K as marker proteins. Control cells and cells with stable knockdown of FMR1 or FXR1 were used. b, Co-IP experiments with or without RNase A in HeLa cells between ADAR1, ADAR2, FMRP and FXR1P. Endogenous proteins were targeted except for ADAR2 (where a FLAG-tagged ADAR2 was expressed). c, Shortest distance between FMRP or FXR1P eCLIP peaks and turquoise editing sites resulted from the WGCNA analysis (orange). Ten sets of random control sites (gray) were constructed for comparison (see Methods). d, Experimental testing of an RNA editing site in the TEAD1 gene for its dependence on ADAR1, ADAR2, FMRP or FXR1P. Control HeLa cells or cells with stable knockdown of one of these proteins were used to express a minigene that contains the editing site. Editing levels were measured by Sanger sequencing. Example sequencing traces are shown for each sample with the targeted editing site underlined. Box plots are based on three biological replicates. P values were calculated by Student’s t-test. *: P < 0.05; **: P < 0.01. e, Similar as d, for an editing site in the EEF2K gene. +: p=0.06. f. similar as d, but displaying editing levels of the TEAD1 editing site in minigenes with the wild-type sequence or different versions of mutants introduced to predicted FMRP binding motifs (see Fig. S8a). Wild-type HeLa cells were used to express these minigenes. g, Similar as f, for the editing site in the EEF2K gene (see Fig. S8b). h, RNA editing levels in control HeLa cells and cells with stable knockdown of ADAR1, ADAR2, FMR1 or FXR1. Hyper-editing sites in three genes were tested. Error bars represent standard errors of three biological replicates.

We next asked whether the Fragile X proteins interact with the ADAR proteins. Reciprocal Co-immunoprecipitation (Co-IP) experiments using HeLa cells showed that FMRP interacts with both ADAR1 and ADAR2 in an RNA-independent manner (Fig. 3b). Note that a FLAG-tagged ADAR2 was expressed in the cells for the IP experiment due to inefficient ADAR2 antibodies. Intriguingly, we also observed RNA-independent interaction between ADAR1 and FXR1P, but not between ADAR2 and FXR1P. Additionally, we observed a positive interaction between FMRP and FXR1P (consistent with previous literature^50^), but not between ADAR1 and ADAR2.

### FMRP and FXR1P binding relative to dysregulated editing sites

To further examine the roles of FMRP and FXR1P in RNA editing regulation, by capturing the global binding patterns of RBPs to RNA transcripts, we carried out enhanced UV crosslinking and immunoprecipitation (eCLIP) experiment^51^ in postmortem frontal cortex from control subjects (Methods). Data of two replicated eCLIP experiments and an input control experiment were obtained for each protein (Fig. S7a).

FMRP and FXR1P eCLIP peaks were identified in each replicate (Methods, Table S5), which demonstrated highly correlated read abundance (Fig. S7b). In the analyses hereafter, we combined the peaks from the duplicate experiments for each protein to maximize the sensitivity of peak detection. The peak binding sites of both proteins were predominantly distributed in 3’ UTRs, introns and exons of known genes (Fig. S7c), consistent with previous literature^48,52^. A sequence motif resembling ACUG was identified as the most enriched motif among the significant FMRP eCLIP peaks (Fig. S7d). Importantly, this motif matches a FMRP binding motif previously reported in the literature^52^. For FXR1P, motif analysis using alternative methods converged to one significant motif with the consensus CAUGC (Fig. S7e). To our knowledge, no explicit FXR1P binding motif is known. Nevertheless, the consensus motif identified here contains an AU, consistent with a previous report that FXR1P tends to associate with AU-rich elements^53^. These consistent results confirmed the quality of our eCLIP experiments.

Next, we examined the FMRP and FXR1P binding peaks relative to dysregulated editing sites in ASD frontal cortex. Remarkably, the FMRP and FXR1P eCLIP peaks were significantly enriched around editing sites in the turquoise module compared to random adenosines in region-matched controls (Fig. 3c, Methods). This result suggests that FMRP and FXR1P proteins may regulate RNA editing directly in ASD.

### FMRP directly modulates RNA editing in vivo

To investigate whether FMRP directly affects RNA editing, we carried out a series of minigene reporter assays to analyze the impact of FMRP on two example editing sites (Table S3). These editing sites, located in the 3’ UTRs of the *TEAD1* and *EEF2K* genes respectively, were chosen because they are in close proximity to putative FMRP binding motifs (Fig. S8). Minigenes containing the editing sites and 500-900 nt of their neighborhood were transfected into control HeLa cells, or HeLa cells with *FMR1*, *ADAR1* or *ADAR2* knockdown (Fig. S9a, Methods).

Knockdown of *FMR1* caused significant reduction of editing level at the *TEAD1* editing site (Fig. 3d). In addition, *ADAR2*, but not *ADAR1*, knockdown also caused nearly complete loss of RNA editing (Fig. 3d). Similarly, knockdown of *FMR1* caused significant reduction of *EEF2K* editing level (Fig. 3e). However, this editing site showed a trend of reduction upon *ADAR1* knockdown (p = 0.06), but not *ADAR2* knockdown. *EEF2K* is also endogenously edited in HeLa cells, and responded to *FMR1* and *ADAR1* knockdown significantly, concordant with the minigene assays (Fig. S9b, c). These data are consistent with our observation that FMRP interacts with both ADAR1 and ADAR2 proteins. Moreover, the positive regulation of FMRP on these editing sites is consistent with the positive association of the turquoise eigen-editing site with *FMR1* expression levels (Fig. 2d).

To examine whether FMRP regulates *TEAD1* and *EEF2K* editing through direct interactions with the RNA, we introduced mutations to the FMRP binding motifs in the minigenes in order to weaken the protein-RNA interaction (Fig. S8). Loss of these FMRP binding sites caused significant reduction in RNA editing (Fig. 3f, g). It should be noted that these mutations did not cause significant changes in the predicted double-stranded RNA (dsRNA) structures in the minigenes (Fig. S8), excluding the possibility that the observed editing change is due to alteration of dsRNA structure. The *TEAD1* and *EEF2K* editing sites are likely site-specific editing sites since no other sites were observed in their immediate neighborhood. Our results suggest that FMRP directly regulate the editing of these two site-specific sites.

### FXR1P regulates hyperedited sites in vivo

In contrast to site-specific editing, another class of editing sites consists of hyperedited sites that tend to cluster together^36^. To examine hyperediting, we carried out minigene experiments on three genes (*CNTNAP4*, *NLGN1*, and *TENM2*), two of which (*CNTNAP4* and *NLGN1*) are ASD candidate risk genes. All three genes have multiple observed editing sites located in long double-stranded intronic regions (Fig. S10, Table S3). *ADAR1* knockdown caused reduction in all detectable editing sites within these long double stranded regions (Fig. 3h), which is consistent with previous literature that ADAR1 regulates promiscuous editing sites^54^. These editing sites also responded to *ADAR2* knockdown, but to a lesser degree compared to their responses to *ADAR1* knockdown. Remarkably, in contrast to the *TEAD1* and *EEF2K* editing sites that depended on FMRP (but not FXR1P, Fig. 3d, e), the hyperedited sites in these genes showed increased editing levels in *FXR1* (but not *FMR1*) knockdown cells. This negative regulation of FXR1P on RNA editing is consistent with the WGCNA results that showed negative correlation between *FXR1* expression and RNA editing (Fig. 2d). Furthermore, RNA immunoprecipitation experiments support that FXR1P binds to the regions harboring the differential editing sites in these target genes (Fig. S11). Together, our results clearly validate that FMRP and FXR1P are important regulators of RNA editing.

### Convergent RNA editing alterations between ASD and Fragile X patients

Loss of FMRP in human leads to the neurodevelopmental disease, Fragile X syndrome, which is also the most frequent monogenic cause of ASD accounting for approximately 1-2% of all ASD cases^31,55^. Additionally, approximately 50% of patients with Fragile X are co-diagnosed with ASD^56^, further highlighting shared molecular underpinnings between ASD and Fragile X syndrome^55^. To understand how these two diseases may share aberrations in RNA editing, we generated RNA-Seq data of the frontal cortex of two patients with Fragile X syndrome and two Fragile X carriers. Western blot confirmed that FMRP expression was absent or reduced in the Fragile X samples relative to carriers, and the expression levels of ADAR1 and ADAR2 are similar between the two groups (Fig. S12a).

RNA editing sites were identified using the RNA-Seq data (Fig. S12b, c) and their editing levels were compared between the Fragile X patients and carriers (Fig. S12d, e, Methods). Strikingly, differential editing sites resulting from this analysis demonstrated a predominant trend of hypoediting in Fragile X patients (Fig. 4a, Table S6), similarly to observations in the idiopathic ASD patients (Fig. 1c). Importantly, these differential editing sites were also more enriched around FMRP and FXR1P eCLIP peaks than expected by chance (Fig. 4b). Moreover, a statistically significant overlap was observed between the differential editing sites in Fragile X patients and those in the turquoise module of ASD frontal cortex (Fig. 4c), the module that is correlated with FMRP expression (Fig.2d). This result again supports our hypothesis that the turquoise module encapsulates a subset of dysregulated editing sites in ASD that are under regulation by FMRP.

**Figure 4.**
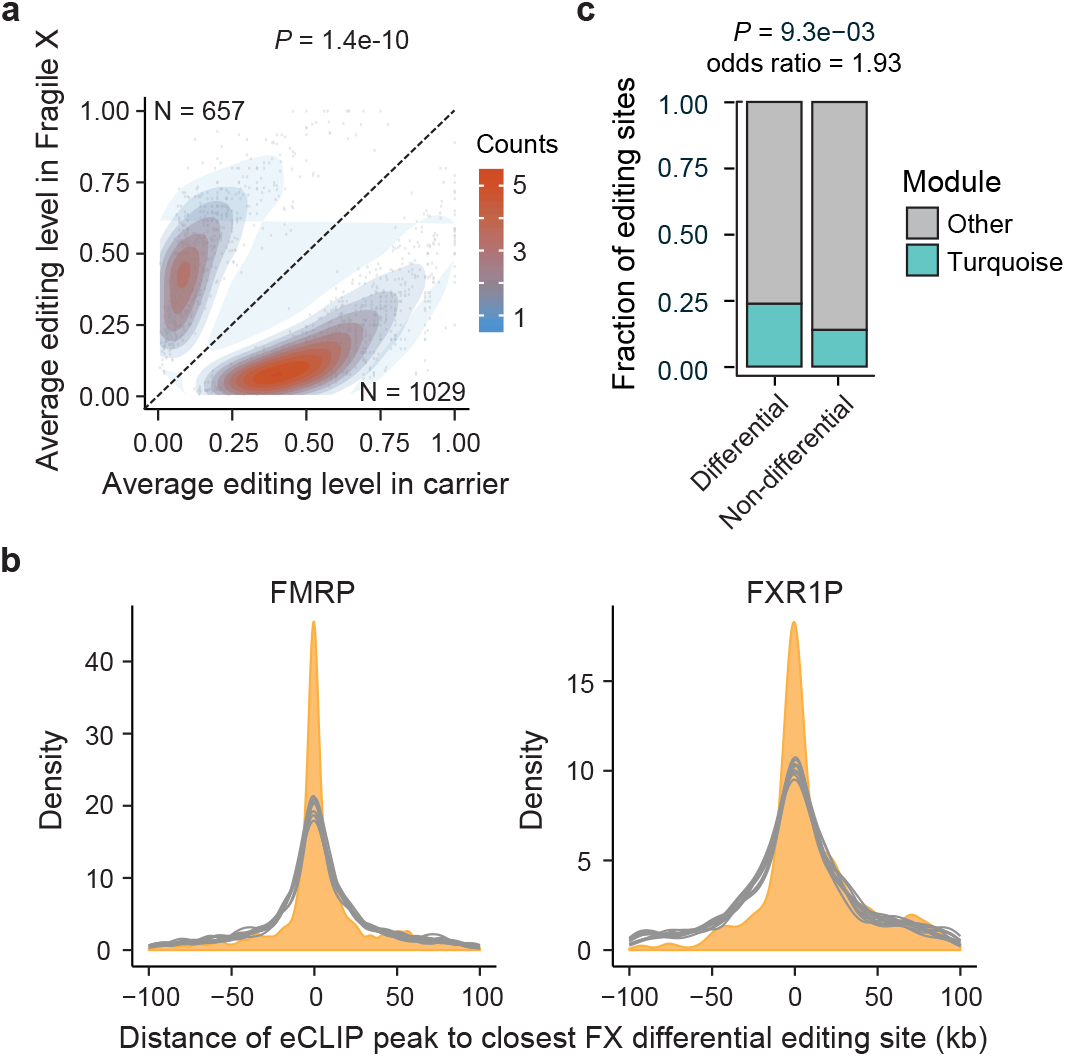
Transcriptome-wide differential editing in the frontal cortex of Fragile X patients and carriers. a, Average editing levels of differential editing sites in Fragile X patients and carriers. Numbers of editing sites that were up- or down-regulated in the patients are shown, which were compared via Fisher’s Exact test (P value shown above plot). b, Shortest distance between FMRP or FXR1P eCLIP peaks and differential editing sites in a (orange). Ten sets of random control sites (gray) were constructed for comparison (see Methods). c, Overlap between the WGCNA turquoise sites of ASD frontal cortex and differential editing sites in the Fragile X patients. P value was calculated by Fisher’s Exact test.

Altogether, the analysis of editing profiles in Fragile X patients provides a strong independent line of evidence showing convergence of dysregulated RNA editing between Fragile X syndrome and ASD through a common mechanism involving FMRP regulation of RNA editing.

### Consistent hypoediting patterns observed for different brain regions of ASD patients

In addition to the frontal cortex, the temporal cortex and cerebellum are also functionally implicated in ASD pathophysiology^57^. Here we ask whether different brain regions have similar editing patterns. Similar to frontal cortex, we observed global down-regulation of RNA editing in both brain regions in ASD (Fig. 5a). In addition, differential editing sites shared between brain regions showed significant correlation in levels of dysregulation (Fig. 5b). We next carried out WGCNA analysis on the editing sites identified in temporal cortex and cerebellum, respectively. Similarly as for frontal cortex, the module (colored turquoise by WGCNA convention) associated with each of these brain regions was also strongly associated with ASD (Fig. 5c, Table S4). Importantly, the turquoise modules of the three brain regions shared many editing sites (Fig. 5d).

**Figure 5.**
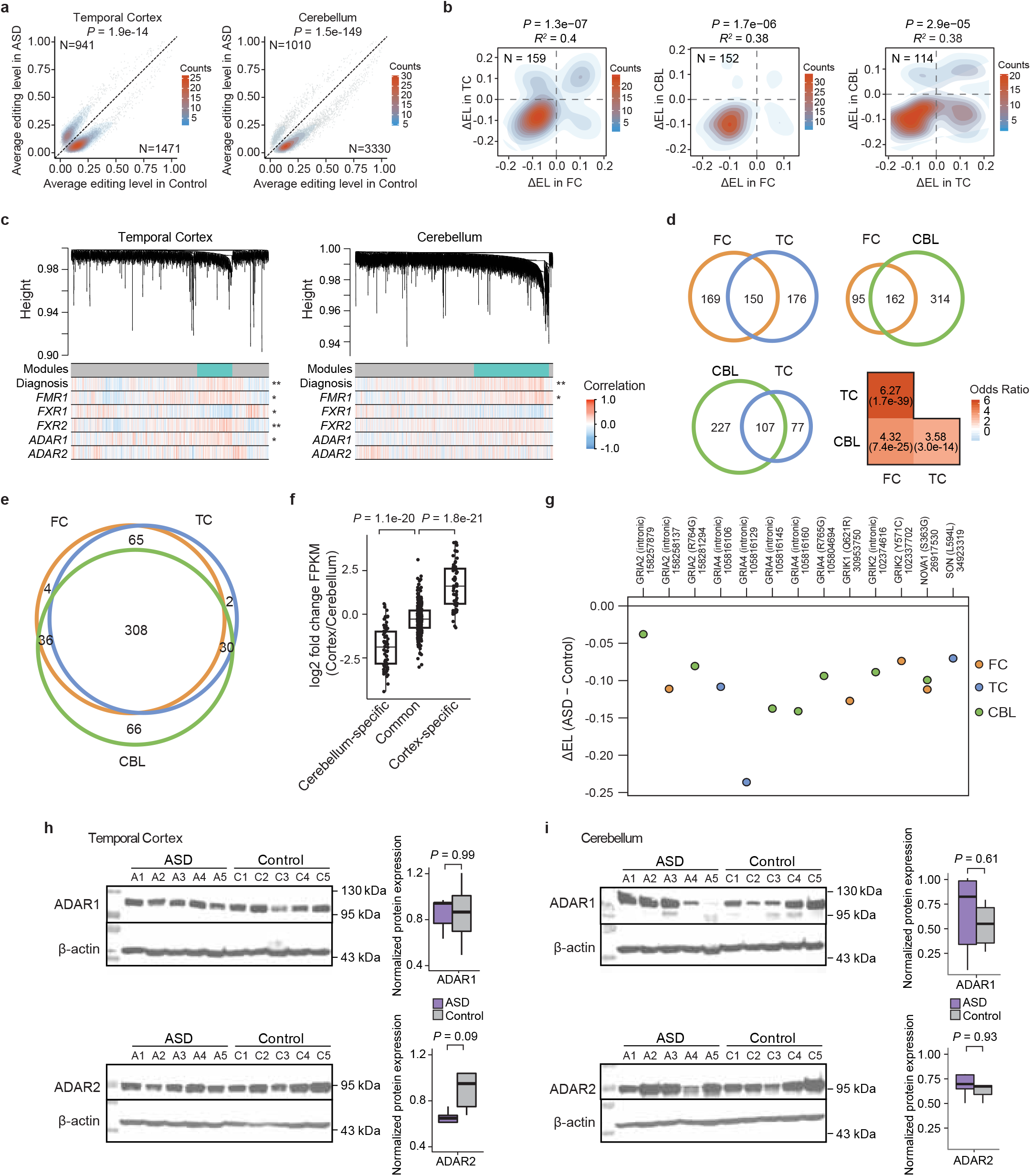
RNA editing dysregulation in different brain regions. a, Average editing levels of differential editing sites in ASD and controls. Similar as Fig. 1c, but data for temporal cortex and cerebellum are shown respectively. b, Changes in editing levels (ΔEL) between ASD and control (ASD-control) of differential editing sites shared across brain regions. TC: temporal cortex; FC: frontal cortex; CBL: cerebellum. Pearson’s correlation p and R^2^ values are shown. c, Similar as Fig. 2d, for WGCNA analysis of editomes in the temporal cortex and cerebellum regions. P values of Pearson’s correlation between diagnosis or mRNA expression levels and eigenediting sites of the turquoise modules were calculated: *: P < 0.05; **: P < 0.01. d, Overlap of editing sites in the turquoise modules of pairs of brain regions. Only editing sites with sufficient read coverage in both brain regions for WGCNA analysis are included. Odds ratios and P values for the overlaps are shown in the heatmap (Fisher’s Exact test). e, Overlap of genes harboring differential editing sites across brain regions. f, Relative FPKM values (log2 fold change) between cortex and cerebellum samples for genes that harbor differential editing sites only in cerebellum, only in cortex or in both types of regions. P values were calculated by Student’s t-test. g, Editing level difference (ΔEL, ASD-control) for a small number of literature-reported evolutionarily conserved RNA editing sites that showed differential editing between ASD and control groups in at least one brain region. h, Similar as Fig. 2b, Western blot of ADAR1 and ADAR2 proteins in temporal cortex samples. Samples used in this experiment are shown in Table S1. i, Similar as h, Western blot of ADAR1 and ADAR2 proteins in cerebellum samples.

Nevertheless, we did observe a small set of 65 and 66 genes exclusively differentially edited in cortex and cerebellum respectively (Fig. 5e, Table S7). These genes exhibited significant cortex- and cerebellum-specific expression patterns (Fig. 5f), suggesting that the region-specific differential editing may be explained by higher expression in their respective brain regions. It is likely that these region-specific genes have distinct functional roles in ASD.

We also examined the editing patterns of 59 editing sites that are known to be conserved across multiple phylogenetic taxa and thus likely represent the most functionally important RNA editing sites in human^58^. Among these editing sites, 13 were identified as differentially edited (p<0.05) in at least one brain region of the ASD patients. Strikingly, all 13 sites demonstrated the hypoediting trend in ASD, 6 of which were recoding sites (Fig. 5g). Four of the recoding sites are located in glutamate receptors, that is, *GRIA2* (R764G), *GRIA4* (R765G), *GRIK1* (Q621R) and *GRIK2* (Y571C)^21^. Additionally, another recoding site was found in the *NOVA1* gene (Fig. 5g), which codes for a brain-specific splicing factor previously reported to cause down-regulated splicing in ASD^59^. This recoding site (S363G) is known to stabilize protein half-life of NOVA1^59^, suggesting that the down-regulated editing may be an upstream causal factor of down-regulated splicing in ASD^15^. Overall these findings strengthen the association between RNA editing and aberrant synaptic signaling in ASD.

### Common and brain region-specific mechanisms of RNA editing regulation in ASD

Finally, we examined the correlation of the turquoise module with expression levels of the *ADAR* and Fragile X-related genes in all three brain regions. Similar to those in frontal cortex, the eigen-editing sites of the turquoise modules in the other two brain regions also displayed correlations with both *FMR1* and *FXR1* expression (Fig. 5c, although the correlation for *FXR1* in cerebellum was not statistically significant, p = 0.07), suggesting that regulation of RNA editing by FMRP and FXR1P may be a common mechanism for multiple afflicted brain regions in ASD.

Although we did not observe significant changes of the *ADAR* mRNAs between ASD and control groups in any brain region (Fig. S13a), we found that *ADAR2* was significantly associated with the overall principal component of differential editing in temporal cortex (Fig. S13b). Interestingly, Western blot analysis of *ADAR* protein levels in temporal cortex and cerebellum also identified a possible trend of *ADAR2* down-regulation in the temporal cortex of ASD (Fig. 5h). Furthermore, *FXR2*, though not associated with the turquoise module in frontal cortex, showed significantly positive correlation with editing in temporal cortex (Fig. 5c). Future studies are needed to examine the functions roles of *FXR2* and *ADAR2* in these brain regions.

## Discussion

Here we perform the first global investigation of RNA editing in ASD and uncover a common trend of hypoediting in ASD patients across different brain regions. Furthermore, we showed that the *FMR1* and *FXR1* genes correlate with the hypoediting patterns in ASD and are direct regulators of RNA editing in human. Consistent with the roles of FMRP and FXR1P in RNA editing regulation, we further demonstrated convergent RNA editing patterns between ASD and Fragile X syndrome, revealing a shared molecular deficit in these closely related neurodevelopmental disorders.

As the monogenic cause of the Fragile X syndrome, FMRP has been the focus of many studies of ASD. For example, genes with rare de novo mutations^32,60,61^, common variation^62^, and copy number variants^63^ in ASD are enriched in FMRP target genes^48^. Multiple transcriptome analyses identified significant correlation between FMRP expression and ASD-associated gene modules^14,33^. Many similar cognitive and behavioral symptoms manifest in both ASD and Fragile X syndrome^55^. In addition, FMRP was shown to be downregulated in ASD patients^64,65^. The plethora of related literature supports the involvement of FMRP in the pathogenesis of ASD and highlights the need to elucidate its potential molecular mechanisms. Our study addresses this question by showing that RNA editing may be an important mechanism via which FMRP contributes to the molecular abnormalities observed in ASD.

Our data supports a model where FMRP interacts with the ADAR proteins to promote editing by directly interacting with the RNA substrates. This model is consistent with previous studies of FMRP in Mouse^66^, Drosophila^67^, and Zebrafish^68^. In contrast, we observed that FXR1P represses editing by interacting with ADAR1 and the RNA targets. The involvement of FXR1P in RNA editing regulation has not been previously reported. Intriguingly, besides the opposite direction in their impact on RNA editing, we found that FMRP and FXR1P showed distinct features among the regulatory targets validated in our study. FXR1P often acted on promiscuous sites and FMRP appeared to regulate site-selective editing sites. It is not yet clear whether FMRP and FXR1P have synergistic functions in the regulation of RNA editing, although these two proteins have shared functions in other biological processes quite relevant to neurodevelopmental disorders, such as neurogenesis^46,69^.

Our study revealed substantial similarities in global editing changes in ASD across the three brain regions we profiled. This highly reproducible editing pattern across regions indicate that dysregulation of RNA editing may affect molecular pathways that are essential to general neurological function. Nevertheless, our data also pointed to possible existence of region-specific editing regulation. For example, we observed a downregulation trend of ADAR2 proteins in the temporal cortex of ASD patients, but not in the frontal cortex or cerebellum. In addition, expression levels of the gene FXR2, an autosomal homolog of FMR1, demonstrated strong correlation with RNA editing levels, which is again a temporal cortex-specific observation (Fig. 5c). These data suggest that future studies aimed at studying region-specific RNA editing will likely uncover novel region-specific regulatory mechanisms.

We observed that RNA editing alterations occurred in genes of critical neurological relevance. Notably, a number of differential editing sites were found in genes involved in synaptic development and homeostasis (Fig. S6), including contactins (*CNTNAP2*^70^, *CNTNAP4*^71^), neurexins (*NRXN1*^72^, *NRXN3*^73^), ankyrins (*ANK2*^32^), and neuronal splicing factors (*NOVA1*^15^ and *RBFOX1*^74^), all of which are known to harbor genetic mutations associated with ASD^42^. These observations indicate that aberrant RNA editing may act together with rare mutations and common genetic perturbations to constitute the pathologic basis of ASD phenotypes. This current work indicates that it will be important to further explore the role RNA editing in ASD pathophysiology, so as to determine whether these changes are causal, or reflect homeostatic or dyshomeostatic responses.

## Methods

### RNA-Seq data sets of ASD and control brain samples

We obtained RNA-Seq data sets of three brain regions of ASD and control subjects from our previous study^15^. We used all data sets except (1) samples from subjects < 7 years old (which showed outlying expression patterns compared to all other samples), and (2) samples containing a 15q duplication, an established genetic cause of syndromic ASD^75^. We confirmed that ASD diagnosis was not confounded by age, batch, and other biological and technical variables (Fig. S1). The final sample set consisted of an approximately equal number of controls and ASD samples totaling 62 samples in frontal cortex, 57 samples in temporal cortex, and 60 samples in cerebellum (Table S1).

### RNA-Seq data sets of Fragile X patients and carriers

Postmortem frontal cortex samples of Fragile X patients and Carriers were obtained from the University of Maryland Brain and Tissue Bank. Total RNA was extracted using TRIzol (Thermo Fisher Scientific, 15596018). RNA-Seq libraries were prepared using NEBNext Poly (A) mRNA magnetic isolation module (NEB, E7490) followed by NEBNext Ultra Directional RNA library prep kit for Illumina according to manufacturer’s instruction. RNA-Seq data were collected on an Illumina HiSeq 2000 sequencer. Paired end reads (2x100nt) were obtained with about 75 million read pairs per sample (Fig. S12b).

### RNA-Seq read mapping and RNA editing identification

RNA-Seq reads were mapped using RASER^76^, an aligner optimized for detecting RNA editing sites, using parameters m = 0.05 and b = 0.03. Uniquely mapped read pairs were retained for further analysis. Unmapped reads were extracted and processed to identify “hyperediting” sites. A recent study showed that previous RNA editing identification methods failed to detect editing sites in hyperedited regions due to existence of a high number of mismatches in the reads^36^.

Our implementation of the hyperediting pipeline closely followed a strategy used by a previous study^36^. In brief, all adenosines in unmapped reads were converted into guanosines. These reads were aligned to a modified hg19 genome where adenosines were also substituted by guanosines. Uniquely mapped read pairs were obtained from this alignment step, and previously converted adenosines were reinstituted. We then combined these hyperedited reads with the originally uniquely mapped reads to identify RNA editing sites.

The procedures described in our previous studies were used to identify RNA editing sites^34,35^. First, RNA-DNA differences (RDDs) were identified as mismatches between the reads and the human reference genome. A log-likelihood test was carried out to determine whether an RDD is likely resulted from a sequence error^34^. A number of posterior filters were then applied to remove RDD sites that were likely caused by technical artifacts in sequencing or read mapping^35^.

Due to limited sequencing depth and the inherent nature of random sampling in RNA-Seq, some editing sites are observable in only a small number of subjects within a population cohort. Editing sites with low apparent prevalence lack sufficient sample size to enable a comparison between ASD and control groups. Therefore, we applied the following filters to retain a subset of editing sites: (1) in each sample, an editing site was required to have at least 5 total reads among which at least 2 reads were edited; (2) the editing site should satisfy filter (1) in at least 5 samples. We applied these filters to editing sites called within each brain region separately.

### Identification of differential RNA editing sites

We define differential RNA editing sites as those: (1) that had significantly different average editing levels between ASD and controls, or (2) that were observed at significantly different population frequencies. A challenge with statistical testing for differential editing levels is that editing level estimation has a larger uncertainty at lower read coverage. More accurate calculations could be obtained by setting a high threshold for read coverage. However, this remedy leads to fewer samples or reduced power per editing site. We developed a strategy that attempts to optimize the trade-off between read coverage requirement and sensitivity in detecting differential editing.

Specifically, the following procedures were implemented for each editing site *e_i_*. (1) we first identified the highest total read coverage requirement for *e_i_* at which there were at least 5 samples in both control and ASD groups. The following read coverage was considered: 20, 15 and 10, in the order of high to low. (2) If a read coverage requirement *C* was reached in (1), we calculate thed average editing level of *e_i_* among the ASD and control samples (*M_ai_*, *M_ci_*), respectively, that satisfied *C*. (3) We then considered samples where *e_i_* did not have at least *C* reads, but satisfied a lower read coverage cutoff (15, 10, or 5). These samples were included if their inclusion did not alter *M_ai_* and *M_ci_* by more than 0.03. (4) We carried out Wilcoxon rank-sum test between editing levels of the above samples in ASD and control groups. (5) If a read coverage requirement *C* was not reached in (1), then we tested all samples where *e_i_* had ≥ 10 read coverage so long as there were at least 10 ASD and 10 control samples. Differential editing sites were defined as those with a p value < 0.05 and an effect size > 5%.

Another type of differential editing was defined as editing sites that have different prevalence between ASD and controls. For each editing site, a Fisher’s Exact test was carried out to compare the numbers of ASD and control samples with nonzero editing levels, with the background being the numbers of ASD and control samples with zero editing level. The minimum read coverage requirement per site was obtained using the same adaptive procedure as described above for the first type of differential editing sites. Differential editing sites were defined as those with p < 0.05 and an effect size > 10%.

### Identification of genes enriched with differential editing

This analysis aims to identify genes that are enriched with differential editing sites. One might consider the top differentially edited genes as those with the largest number of differential editing sites. However, as expected, there exists a positive correlation between gene length and the number of differential editing sites (Fig. S6a). Therefore, we used a linear model to construct a regression between these two variables. We defined genes as enriched with differential editing if they had more differential editing sites than expected (beyond 95% confidence interval of the expected mean).

### Principal components analysis (PCA)

PCA was conducted on differential editing sites in order to examine associations between PCs and potential confounding covariates. The R function prcomp was used for this purpose. Missing values in the editing level matrix were imputed using the missMDA package^77^. The PCs were then correlated against technical and biological covariates such as age and gender (Fig. S5a). The first PC was predominantly associated with ASD diagnosis, and was thus used as the PC for differential editing.

### Weighted gene co-expression network analysis (WGCNA)

The WGCNA package^43^ in R was used to find modules of correlated editing sites. In multi-sample analysis, it is typical that some editing sites have no available values (missing data) in certain samples that lack read coverage at those sites. To preclude inaccurate calculations due to samples with too much missing data, we used the following requirements for editing sites to be included in WGCNA: (1) with ≥5 reads in ≥ 90% of samples and (2) with nonzero editing levels in ≥ 10% samples. In addition, to detect variation in the data, we further required that the included editing sites had a standard deviation ≥ 0.1 in their editing levels across samples. A soft threshold power of 10 was used to fit scale-free topology. To avoid obtaining modules driven by outlier samples, we followed our previous bootstrapping strategy^15,18^. One hundred bootstraps of the data set were carried out to compute the topological overlap matrix of each resampled network. Co-editing modules were obtained using the consensus topological overlap matrix of the 100 bootstraps.

WGCNA offers a dynamic tree-cutting algorithm, which enables identification of modules at various dendrogram heights and allows delineation of nested modules^78^. However, upon examination of the WGCNA dendrogram (Fig. 2d), we observed only one pronounced module of editing sites. Furthermore, most modules, identified through dynamic tree cutting, were generally unstable, highly dependent on tree cutting parameters. Therefore, we used the traditional constant height tree cutting, provided by WGCNA as cutreeStaticColor, with cutHeight set to 0.9965, which produced the single turquoise module. This is the largest module that is most likely biologically relevant and technically robust. In addition, this module is conserved across brain regions (Fig. 5c).

### Association of modules with ASD diagnosis and RNA binding proteins

To test the association of the turquoise module with diagnosis, we first defined eigen-editing sites as the first principal component of the module, according to WGCNA recommendation^44^. A linear regression model was constructed between the eigen-editing sites and diagnosis, in addition to biological and technical covariates including RIN, age, gender, sequencing batch, PMI, brain bank, 5’ to 3’ RNA bias, AT dropout rate, GC dropout rate, mapped bases in intergenic regions, uniquely mapped reads. The linear model was fit with backwards selection, and the module was deemed as associated with ASD diagnosis if p ≤ 0.05 for the coefficient of this variable.

We tested if a module was enriched with differential editing sites using Fisher’s Exact test. In addition, we tested the association between modules and potential regulatory genes by examining the correlation between the eigen-editing sites and mRNA expression levels of the genes. It should be noted that the mRNA expression levels were corrected values following removal of variability contributed by technical covariates^15^.

### eCLIP-Seq experiment and data analysis

The eCLIP experimental procedure is detailed in our previous studies^51,79^. The antibodies used for this experiment are: FMRP antibody (MBL, RN016P) and FXR1 antibody (Bethyl Laboratories, A303-892A). Flash-frozen brain tissue was cryo-ground in pestles pre-chilled with liquid nitrogen, spread out in standard tissue culture plates pre-chilled to −80°C, and UV crosslinked twice at 254 nM (400 mJ/cm^2^). 50 mg of crosslinked tissue was then used for each eCLIP experiment, performed as previously described^51,79^.

eCLIP-Seq data were analyzed using the CLIPper software^51^ that generated a list of predicted binding peaks of the corresponding protein. In each replicate, peaks were further filtered to retain those whose abundance was at least 2 fold of that in the SMInput sample. To examine the FMPR or FXR1P binding relative to RNA editing sites, we compared the distances from eCLIP peaks to turquoise editing sites compared to region-matched random adenosines. Only editing sites residing in genes containing at least 1 eCLIP peak were considered. The closest distance between an editing site/random adenosine and eCLIP peaks were calculated, and density plots were generated using the geom_density function in the ggplot2 package in R. A total of 10 sets of randomly selected control adenosines were generated.

To identify the motifs enriched in eCLIP peaks, we used two alternative methods: HOMER^80^, and DREME^81^. We ran DREME with all eCLIP peaks of each protein using default parameters, which creates control sequences through dinucleotide shuffling. HOMER was run with the findMotifsGenome.pl program (-p 4 -rna -S 10 -len 5,6,7,8,9). Background controls were defined as randomly chosen sequences in the same type of genic region as the true peaks. The control sequences have one-to-one match in length with the actual peaks. Three sets of random controls were constructed. Homopolymer or dinucleotide repeats were discarded. We required the final consensus motif to be the most enriched motif identified by HOMER that was also one of the most enriched motifs resulting from DREME.

### RNA editing analysis of Fragile X samples

The RNA-Seq data derived from Fragile X patients and carriers were analyzed similarly as those of the ASD cohorts. Fisher’s Exact test was used to identify differentially edited sites using pooled patient and carrier data sets (p ≤ 0.05 and effect size > 5%).

### Gene ontology enrichment analysis

Gene ontology (GO) terms were downloaded from Ensembl^82^. For each query gene, a random control gene was chosen to match gene expression level and gene length (±10% relative to that of query gene). GO terms present in the sets of query genes and control genes were collected respectively. A total of 10,000 sets of control genes were obtained. For each GO term, a Gaussian distribution was fit to the number of control genes containing this GO term. The enrichment p value of the GO term among the query genes was calculated using this distribution.

### Validation of RNA editing levels

#### RNA Extraction

Brain tissues were homogenized in RNA TRIzol reagent (Thermo Fisher Scientific, 15596018). Mixture was incubated on ice for 15 min. Chloroform was added to the mixture and incubated at room temperature for 10 min. The mixture was centrifuged at 12000g for 15 min, and the top layer was carefully extracted. Equal volume of 200-proof ethanol was added to the top chloroform layer and mixed thoroughly. RNA was further purified using Direct-zol™ RNA MiniPrep Plus kit (Zymo Research, R2072) following the manufacture’s protocol.

#### cDNA synthesis and PCR

cDNA synthesis was carried out using random hexamers, 1 μg total RNA, and the SuperScript IV Reverse Transcriptase (Thermo Fisher Scientific, 18090050) as described in the manufacturer’s protocol. Next, 2uL cDNA (corresponding to 1/10^th^ of the original RNA) was used as template for PCR reactions using the DreamTaq PCR Master Mix (2X) (Thermo Fisher Scientific, k1082). PCR was performed on an Eppendorf thermal cycler using the following thermal cycle conditions for all candidate sites (5 min, 95°C for hot start followed by 30 cycles of 15 s, 95°C; 15 s, 55°C and 1min/kb, 72°C).

#### Topo Cloning and Clonal Sequencing

PCR products were run on 1% agarose gel and visualized under UV light. The correct size band was cut and digested by Zymoclean™ Gel DNA Recovery Kit (Zymo Research, D4002) following the manufacturer’s protocol. PCR product was inserted into kanamycin resistant pCR 2.1-TOPO vector (Thermo Fisher Scientific, 450641). Ligated clones were transformed into One Shot TOP10 Chemically Competent E. coli (Thermo Fisher Scientific, C404003). Transformed cells were streaked on LB/Agar plates containing kanamycin and X-Gal as selection markers and incubated overnight at 37°C. Each plate was divided into 4 quadrants and 6 white clones were randomly selected from each quadrant (total of 24 clones per patient sample per editing site). Each clone was inoculated overnight in LB/Kanamycin. Plasmid was extracted using Plasmid DNA Miniprep Kits (Thermo Fisher Scientific, K210011). Miniprep samples were sequenced using Genewiz Sanger sequencing. The number of the clones presenting G peak at the editing site of interest was counted to determine the estimated editing ratio.

### Co-immunoprecipitation

Hela Cells were maintained with DMEM supplemented with 10% FBS and 100 U ml-1 penicillin/ streptomycin at 37 °C and 5% CO2. Ten million HeLa cells were collected and lysed in 1 ml non-denaturing lysis buffer at pH 8.0, containing 20 mM Tris-HCl, 137 mM NaCl, 1% NP-40, and 2 mM EDTA supplemented with complete protease inhibitor cocktail. Extracted proteins were incubated overnight with ADAR1 antibody (Santa Cruz, sc-271854) or FMRP antibody (Millipore, MAB2160) at 4 °C; precipitation of the immune complexes was performed with Dynabeads Protein G (Thermo Fisher Scientific, 1003D) for 4 h at 4 °C, according to the manufacturer’s instructions. For experiments involving Flag-ADAR2, the supernatant derived from Flag-tagged hADAR2 overexpressing cells was incubated for 3 h at 4 °C with Flag M2 antibody (Sigma, F1804). After immunoprecipitation, the beads were washed three times with the lysis buffer at 4 °C, and eluted from the Dynabeads using elute buffer (0.2 M glycine, at pH 2.8). Twenty microliters were loaded onto the gel and the samples were processed by SDS-polyacrylamide gel electrophoresis (SDS-PAGE) and analyzed by Western blot. The following antibodies were used for the Western blots: ADAR1 antibody (Santa Cruz, sc-73408), Flag antibody (sc-807), FMRP antibodies (Millipore, MAB2160 and Abcam, ab17722), FXR1P antibody (Bethyl Laboratories, A303-892A), and FXR2 antibody (Sigma-Aldrich, F1554). The HRP-linked secondary antibodies were used and the blots were visualized with the ECL kit (GE, RPN2232).

### Subcellular fractionation

Cells were fractionated following a previously published protocol with some modifications^83^. Briefly, monolayers of cells in 10-cm plates were washed twice with ice-cold PBS, followed by gentle scraping of cells. Cells were resuspended with the ice-cold HLB+N buffer (10 mM Tris-HCl, at pH 7.5, 10 mM NaCl, 2.5 mM MgCl2 and 0.5% NP-40) on ice for 5 min. Lysates were layered over a chilled 10% sucrose cushion made in the ice-cold HLB+NS buffer (10 mM Tris-HCl, at pH 7.5, 10 mM NaCl, 2.5 mM MgCl2, 0.5% NP-40 and 10% sucrose) and centrifuged for 5 min at 4 °C at 420g. After centrifugation, the supernatant was collected and served as the cytoplasmic fraction. The nuclear pellet was then treated with the ice-cold nuclei lysis buffer (10 mM HEPES, at pH 7.6, 300 mM NaCl, 7.5 mM MgCl2, 0.2 mM EDTA, 1 mM DTT, 1 M Urea, and 1% NP-40) after washing. Fractionation efficiency was validated by Western blot using antibody specific to the marker for each fraction: β-tubulin (Sigma, T8328) for the cytoplasmic fraction and rabbit polyclonal U1-70k (a kind gift from Dr. Douglas Black) for the nucleoplasmic fraction.

### Construction of minigenes and site-directed mutagenesis

Partial 3’ UTRs of EEF2K and TEAD1 were restriction digested and inserted between the SacII/XhoI sites in the pEGFP-C1 vector. Overlapping oligonucleotide primers containing the desired mutations were used to amplify mutation-containing fragments from the wild-type minigene plasmid, using Q5 High-Fidelity DNA polymerase (New England Biolabs, M0491L). All resulting amplification products were confirmed by sequencing.

### Transfection, RNA isolation, RT-PCR amplification, and analysis of RNA editing

HeLa cells were grown on 6-well plates under standard conditions at 37 °C with 5% CO2. Cells were grown to 70% confluence, and transfection was performed using Lipofectamine 3000 (Thermo Fisher Scientific, L3000015) with 100 ng of minigene plasmid. Cells were harvested after 24 h. Total RNA was extracted using TRIzol reagent (Thermo Fisher Scientific, 15596018), followed by treatment with 1 U of DNase I (Zymo Research, E1011-A). RNA was further purified using Direct-zol RNA MiniPrep kit following the manufacture’s instruction (Zymo Research, R2072). Reverse transcription (RT) was performed on 1 μg total RNA for 1 h at 42 °C using random hexamer primer and SuperScript IV (Thermo Fisher Scientific, 18090050). The cDNA products derived from the expressed minigenes were detected by PCR using the pEGFP-C1-specific forward primer and a gene-specific reverse primer. Amplification was performed for 30 cycles, consisting of 30 s at 95 °C, 30 s at 55 °C, and 2 minutes at 72 °C. The products from RT-PCR were resolved on 0.8% agarose gels. The appropriate PCR product was excised and the DNA was extracted, purified, and analyzed by Sanger sequencing. A-to-I editing levels were calculated as relative peak heights (that is, ratio between the G peak height and the combined height of A and G peaks, height G / height A + height G).

### Production of lentivirus and cell transduction for protein knockdown

pLKO1 non-target control-shRNA (SHC016), FMR1-targeting shRNA (TRCN0000059758) or FXR1-targeting shRNA (TRCN0000159153) constructs were used. We produced lentiviruses via cotransfection of pCMV-d8.91, pVSV-G and pLKO1 into HEK293T cells using Lipofectamine 3000 (Thermo Fisher Scientific, L3000015). Transduction was carried out according to the standard protocol of the ENCODE consortium^84^. Briefly, viruses were collected from conditioned media after 48 h co-transfection. Lentivirus-containing media was mixed with the same volume of DMEM media that contain polybrene (8 μg/ml), which was used to infect HeLa cells. After 24 h, cells were incubated with puromycin (2 μg/ml) for 3-7 days. Knockdown efficiency was evaluated by Western blot. Cells were lysed in RIPA containing complete protease inhibitor cocktail. Cell lysates were then resolved through 8% SDS-PAGE and probed by ADAR1 antibody (Santa Cruz, sc-271854), ADAR2 antibody (Santa Cruz, sc-73409), FMRP antibody (Millipore, MAB2160), FXR1P antibody (Bethyl Laboratories, A303-892A), and FXR2 antibody (Sigma-Aldrich, F1554).

### Western Blot in ASD and Fragile-X brain samples

Brain tissues were homogenized in RIPA lysis and extraction buffer containing protease inhibitor (Thermo Scientific, 88866). Mixture was then incubated on ice for 30 minutes, sonicated, and spun down. Crude protein concentration was obtained using Pierce BCA Protein Assay Kit (Thermo Fisher Scientific, 23225). Equal amount of protein was separated using 8% SDS–PAGE and then transferred onto nitrocellulose membrane. The membrane was blocked with 5% non-fat milk (Genesee Scientific, 20-241) and 0.1% Tween 20 in tris-buffered saline. The blot was incubated in primary antibody solution against the protein of interest with 5% non-fat milk and 0.1% Tween 20 in TBS overnight at 4°C on shaker. Antibodies used in this experiment include ADAR1 antibody (Santa Cruz, sc-271854), ADAR2 antibody (Santa Cruz, sc-73409), FMRP antibody (Millipore, MAB2160). Secondary antibody containing goat anti-mouse IgG-HRP (sc-2005, Santa Cruz Biotechnology) was used to label the corresponding primary antibody. The blot was developed using Amersham ECL Prime Western Blotting Detection Reagent (GE Healthcare Life Sciences, RNP2232) and imaged with the Syngene PXi immunoblot imaging system. Beta Actin was used as a loading control. Western blot images were analyzed using ImageJ.

### RNA immunoprecipitation (RIP)-PCR

RIP was performed according to previously published protocols with some modifications^85^. Cells were harvested on the second day of minigene transfection in RIP buffer (25 mM Tris-HCl, at pH 7.4, 150 mM KCl, 5 mM EDTA, 0.5% NP-40 and 0.5 mM DTT supplemented with complete protease inhibitor cocktail and 100 U ml-1 RNase OUT), sonicated (10 s three times with 1 min intervals) and centrifuged at 13,000 rpm for 10 min at 4 °C. Supernatant was treated with 100 U RNase-free DNase I (Zymo Research, E1011-A) at 37 °C for 30 min and then centrifuged again at 13,000 rpm for 10 min at 4 °C. For immunoprecipitation, lysates were incubated with FXR1P antibody (Santa Cruz, sc-374148) or anti-mouse IgG (Santa Cruz, sc-2025) as a negative control overnight at 4 °C. The Dynabeads were washed three times with the RIP buffer and bound RNA was isolated using TRIzol (Thermo Fisher Scientific, 15596018), according to the manufacturer’s instructions. Eluted RNA was reverse-transcribed using SuperScript IV (Thermo Fisher Scientific, 18090050) with random hexamer primers. PCR was carried out for 30 cycles, consisting of 30 s at 95 °C, 30 s at 55 °C, and 30 s at 72 °C. PCR products were analyzed by agarose gel electrophoresis.

## Data availability

All sequencing data (eCLIP-Seq of FMRP, FXR1P, RNA-Seq of Fragile-X patients and carriers) are being deposited to GEO (accession number pending). RNA-Seq data sets of ASD and control brains were obtained from our previous study^15^ and are available in the PsychENCODE website (https://www.synapse.org//#!Synapse:syn4921369/wiki/235539).

## Acknowledgements

Postmortem brain samples of used in this study were obtained from the University of Maryland Brain and Tissue Bank, which is a component of the NIH NeuroBioBank. We are grateful to the patients and families who participate in the tissue donation programs. This work was funded by grants from the National Institute of Health to X.X. (HG009417, HG006264), G.W.Y. (HG004659, HG007005 and NS075449), S.T. (T32HG002536), E.L.V.N. (HG009530). S.T is supported by the UCLA Eureka Scholarship. E.L.V.N. is a Merck Fellow of the Damon Runyon Cancer Research Foundation (DRG-2172-13). G.A.P. is supported by the National Science Foundation Graduate Research Fellowship.

## References

1 Association, A. P. Diagnostic and statistical manual of mental disorders (4th ed., text rev.). 4th edn, (2000).

2 Rojas, D. C. The role of glutamate and its receptors in autism and the use of glutamate receptor antagonists in treatment. Journal of neural transmission (Vienna, Austria: 1996) 121, 891–905, doi:10.1007/s00702-014-1216-0 (2014).

3 Yang, P. & Chang, C. L. Glutamate-mediated signaling and autism spectrum disorders: emerging treatment targets. Current pharmaceutical design 20, 5186–5193 (2014).

4 De Grandis, E. et al. Cerebrospinal fluid alterations of the serotonin product, 5-hydroxyindolacetic acid, in neurological disorders. Journal of inherited metabolic disease 33, 803–809, doi:10.1007/s10545-010-9200-9 (2010).

5 Guo, Y. P. & Commons, K. G. Serotonin neuron abnormalities in the BTBR mouse model of autism. Autism research: official journal of the International Society for Autism Research 10, 66–77, doi:10.1002/aur.1665 (2017).

6 Chugani, D. C. Role of altered brain serotonin mechanisms in autism. Mol Psychiatry 7 Suppl 2, S16–17, doi:10.1038/sj.mp.4001167 (2002).

7 Ha, S., Sohn, I. J., Kim, N., Sim, H. J. & Cheon, K. A. Characteristics of Brains in Autism Spectrum Disorder: Structure, Function and Connectivity across the Lifespan. Experimental neurobiology 24, 273–284, doi:10.5607/en.2015.24.4.273 (2015).

8 Nelson, S. B. & Valakh, V. Excitatory/Inhibitory Balance and Circuit Homeostasis in Autism Spectrum Disorders. Neuron 87, 684–698, doi:10.1016/j.neuron.2015.07.033 (2015).

9 Marchetto, M. C. et al. Altered proliferation and networks in neural cells derived from idiopathic autistic individuals. Mol Psychiatry, doi:10.1038/mp.2016.95 (2016).

10 Mariani, J. et al. FOXG1-Dependent Dysregulation of GABA/Glutamate Neuron Differentiation in Autism Spectrum Disorders. Cell 162, 375–390, doi:10.1016/j.cell.2015.06.034 (2015).

11 Geschwind, D. H. & State, M. W. Gene hunting in autism spectrum disorder: on the path to precision medicine. The Lancet. Neurology 14, 1109–1120, doi:10.1016/s1474-4422(15)00044-7 (2015).

12 Abrahams, B. S. & Geschwind, D. H. Advances in autism genetics: on the threshold of a new neurobiology. Nature reviews. Genetics 9, 341–355, doi:10.1038/nrg2346 (2008).

13 Voineagu, I. et al. Transcriptomic analysis of autistic brain reveals convergent molecular pathology. Nature 474, 380–384, doi:10.1038/nature10110 (2011).

14 Gupta, S. et al. Transcriptome analysis reveals dysregulation of innate immune response genes and neuronal activity-dependent genes in autism. Nat Commun 5, 5748, doi:10.1038/ncomms6748 (2014).

15 Parikshak, N. N. et al. Genome-wide changes in lncRNA, splicing, and regional gene expression patterns in autism. Nature 540, 423–427, doi:10.1038/nature20612 (2016).

16 Irimia, M. et al. A highly conserved program of neuronal microexons is misregulated in autistic brains. Cell 159, 1511–1523, doi:10.1016/j.cell.2014.11.035 (2014).

17 Quesnel-Vallieres, M. et al. Misregulation of an Activity-Dependent Splicing Network as a Common Mechanism Underlying Autism Spectrum Disorders. Mol Cell 64, 1023–1034, doi:10.1016/j.molcel.2016.11.033 (2016).

18 Wu, Y. E., Parikshak, N. N., Belgard, T. G. & Geschwind, D. H. Genome-wide, integrative analysis implicates microRNA dysregulation in autism spectrum disorder. Nat Neurosci 19, 1463–1476, doi:10.1038/nn.4373 (2016).

19 Nishikura, K. Functions and regulation of RNA editing by ADAR deaminases. Annual review of biochemistry 79, 321–349, doi:10.1146/annurev-biochem-060208-105251 (2010).

20 Bass, B. L. RNA editing by adenosine deaminases that act on RNA. Annual review of biochemistry 71, 817–846, doi:10.1146/annurev.biochem.71.110601.135501 (2002).

21 Behm, M. & Ohman, M. RNA Editing: A Contributor to Neuronal Dynamics in the Mammalian Brain. Trends Genet 32, 165–175, doi:10.1016/j.tig.2015.12.005 (2016).

22 Nishikura, K. A-to-I editing of coding and non-coding RNAs by ADARs. Nature reviews. Molecular cell biology 17, 83–96, doi:10.1038/nrm.2015.4 (2016).

23 Chen, L. et al. Recoding RNA editing of AZIN1 predisposes to hepatocellular carcinoma. Nat Med 19, 209–216, doi:10.1038/nm.3043 (2013).

24 Sodhi, M. S., Burnet, P. W., Makoff, A. J., Kerwin, R. W. & Harrison, P. J. RNA editing of the 5-HT(2C) receptor is reduced in schizophrenia. Mol Psychiatry 6, 373–379, doi:10.1038/sj.mp.4000920 (2001).

25 Lyddon, R., Dwork, A. J., Keddache, M., Siever, L. J. & Dracheva, S. Serotonin 2c receptor RNA editing in major depression and suicide. The world journal of biological psychiatry: the official journal of the World Federation of Societies of Biological Psychiatry 14, 590-601, doi:10.3109/15622975.2011.630406 (2013).

26 Khermesh, K. et al. Reduced levels of protein recoding by A-to-I RNA editing in Alzheimer’s disease. RNA (New York, N.Y.) 22, 290–302, doi:10.1261/rna.054627.115 (2016).

27 Gaisler-Salomon, I. et al. Hippocampus-specific deficiency in RNA editing of GluA2 in Alzheimer’s disease. Neurobiology of aging 35, 1785–1791, doi:10.1016/j.neurobiolaging.2014.02.018 (2014).

28 Hideyama, T. et al. Profound downregulation of the RNA editing enzyme ADAR2 in ALS spinal motor neurons. Neurobiology of disease 45, 1121–1128, doi:10.1016/j.nbd.2011.12.033 (2012).

29 Kwak, S. & Kawahara, Y. Deficient RNA editing of GluR2 and neuronal death in amyotropic lateral sclerosis. Journal of molecular medicine (Berlin, Germany) 83, 110-120, doi:10.1007/s00109-004-0599-z (2005).

30 Eran, A. et al. Comparative RNA editing in autistic and neurotypical cerebella. Mol Psychiatry 18, 1041–1048, doi:10.1038/mp.2012.118 (2013).

31 de la Torre-Ubieta, L., Won, H., Stein, J. L. & Geschwind, D. H. Advancing the understanding of autism disease mechanisms through genetics. Nature medicine 22, 345–361 (2016).

32 lossifov, I. et al. De novo gene disruptions in children on the autistic spectrum. Neuron 74, 285–299, doi:10.1016/j.neuron.2012.04.009 (2012).

33 Parikshak, N. N. et al. Integrative functional genomic analyses implicate specific molecular pathways and circuits in autism. Cell 155, 1008–1021, doi:10.1016/j.cell.2013.10.031 (2013).

34 Bahn, J. H. et al. Accurate identification of A-to-I RNA editing in human by transcriptome sequencing. Genome research 22, 142–150, doi:10.1101/gr.124107.111 (2012).

35 Lee, J. H., Ang, J. K. & Xiao, X. Analysis and design of RNA sequencing experiments for identifying RNA editing and other single-nucleotide variants. RNA (New York, N.Y.) 19, 725–732, doi:10.1261/rna.037903.112 (2013).

36 Porath, H. T., Carmi, S. & Levanon, E. Y. A genome-wide map of hyper-edited RNA reveals numerous new sites. Nat Commun 5, 4726, doi:10.1038/ncomms5726 (2014).

37 Picardi, E., D’Erchia, A. M., Lo Giudice, C. & Pesole, G. REDIportal: a comprehensive database of A-to-I RNA editing events in humans. Nucleic acids research 45, D750–D757, doi:10.1093/nar/gkw767 (2017).

38 Levanon, E. Y. et al. Systematic identification of abundant A-to-I editing sites in the human transcriptome. Nature biotechnology 22, 1001–1005, doi:10.1038/nbt996 (2004).

39 Kleinberger, Y. & Eisenberg, E. Large-scale analysis of structural, sequence and thermodynamic characteristics of A-to-I RNA editing sites in human Alu repeats. BMC genomics 11, 453, doi:10.1186/1471-2164-11-453 (2010).

40 Hwang, T. et al. Dynamic regulation of RNA editing in human brain development and disease. Nat Neurosci 19, 1093–1099, doi:10.1038/nn.4337 (2016).

41 Rakic, P. Evolution of the neocortex: a perspective from developmental biology. Nature reviews. Neuroscience 10, 724–735, doi:10.1038/nrn2719 (2009).

42 Abrahams, B. S. et al. SFARI Gene 2.0: a community-driven knowledgebase for the autism spectrum disorders (ASDs). Molecular autism 4, 36, doi:10.1186/2040-2392-4-36 (2013).

43 Langfelder, P. & Horvath, S. WGCNA: an R package for weighted correlation network analysis. BMC bioinformatics 9, 559, doi:10.1186/1471-2105-9-559 (2008).

44 Langfelder, P. & Horvath, S. Eigengene networks for studying the relationships between co-expression modules. BMC systems biology 1, 54, doi:10.1186/1752-0509-1-54 (2007).

45 Till, S. M. The developmental roles of FMRP. Biochemical Society transactions 38, 507-510, doi:10.1042/bst0380507 (2010).

46 Davis, J. K. & Broadie, K. Multifarious Functions of the Fragile X Mental Retardation Protein. Trends in genetics: TIG, doi:10.1016/j.tig.2017.07.008 (2017).

47 Castren, M., Haapasalo, A., Oostra, B. A. & Castren, E. Subcellular localization of fragile X mental retardation protein with the I304N mutation in the RNA-binding domain in cultured hippocampal neurons. Cellular and molecular neurobiology 21, 29–38 (2001).

48 Darnell, J. C. et al. FMRP stalls ribosomal translocation on mRNAs linked to synaptic function and autism. Cell 146, 247–261, doi:10.1016/j.cell.2011.06.013 (2011).

49 Kim, M., Bellini, M. & Ceman, S. Fragile X mental retardation protein FMRP binds mRNAs in the nucleus. Mol Cell Biol 29, 214–228, doi:10.1128/mcb.01377-08 (2009).

50 Zhang, Y. et al. The fragile X mental retardation syndrome protein interacts with novel homologs FXR1 and FXR2. The EMBO journal 14, 5358–5366 (1995).

51 Van Nostrand, E. L. et al. Robust transcriptome-wide discovery of RNA-binding protein binding sites with enhanced CLIP (eCLIP). Nature methods 13, 508–514, doi:10.1038/nmeth.3810 (2016).

52 Ascano, M., Jr. et al. FMRP targets distinct mRNA sequence elements to regulate protein expression. Nature 492, 382–386, doi:10.1038/nature11737 (2012).

53 Vasudevan, S. & Steitz, J. A. AU-rich-element-mediated upregulation of translation by FXR1 and Argonaute 2. Cell 128, 1105–1118, doi:10.1016/j.cell.2007.01.038 (2007).

54 Tan, M. H. et al. Dynamic landscape and regulation of RNA editing in mammals. Nature 550, 249–254, doi:10.1038/nature24041 (2017).

55 Hagerman, R., Hoem, G. & Hagerman, P. Fragile X and autism: Intertwined at the molecular level leading to targeted treatments. Molecular autism 1, 12, doi:10.1186/2040-2392-1-12 (2010).

56 Abbeduto, L., McDuffie, A. & Thurman, A. J. The fragile X syndrome-autism comorbidity: what do we really know? Front Genet 5, 355, doi:10.3389/fgene.2014.00355 (2014).

57 Igelstrom, K. M., Webb, T. W. & Graziano, M. S. A. Functional Connectivity Between the Temporoparietal Cortex and Cerebellum in Autism Spectrum Disorder. Cerebral cortex (New York, N.Y.: 1991) 27, 2617–2627, doi:10.1093/cercor/bhw079 (2017).

58 Pinto, Y., Cohen, H. Y. & Levanon, E. Y. Mammalian conserved ADAR targets comprise only a small fragment of the human editosome. Genome Biol 15, R5, doi:10.1186/gb-2014-15-1-r5 (2014).

59 Irimia, M. et al. Evolutionarily conserved A-to-I editing increases protein stability of the alternative splicing factor Nova1. RNA biology 9, 12–21, doi:10.4161/rna.9.1.18387 (2012).

60 Iossifov, I. et al. The contribution of de novo coding mutations to autism spectrum disorder. Nature 515, 216–221, doi:10.1038/nature13908 (2014).

61 De Rubeis, S. et al. Synaptic, transcriptional and chromatin genes disrupted in autism. Nature 515, 209–215, doi:10.1038/nature13772 (2014).

62 Jansen, A. et al. Gene-set analysis shows association between FMRP targets and autism spectrum disorder. European journal of human genetics: EJHG 25, 863–868, doi:10.1038/ejhg.2017.55 (2017).

63 Pinto, D. et al. Convergence of genes and cellular pathways dysregulated in autism spectrum disorders. American journal of human genetics 94, 677–694, doi:10.1016/j.ajhg.2014.03.018 (2014).

64 Fatemi, S. H. & Folsom, T. D. Dysregulation of fragile x mental retardation protein and metabotropic glutamate receptor 5 in superior frontal cortex of individuals with autism: a postmortem brain study. Molecular autism 2, 6, doi:10.1186/2040-2392-2-6 (2011).

65 Fatemi, S. H., Folsom, T. D., Kneeland, R. E. & Liesch, S. B. Metabotropic glutamate receptor 5 upregulation in children with autism is associated with underexpression of both Fragile X mental retardation protein and GABAA receptor beta 3 in adults with autism. Anatomical record (Hoboken, N.J.: 2007) 294, 1635–1645, doi:10.1002/ar.21299 (2011).

66 Filippini, A. et al. Absence of the Fragile X Mental Retardation Protein results in defects of RNA editing of neuronal mRNAs in mouse. RNA biology, 1–12, doi:10.1080/15476286.2017.1338232 (2017).

67 Bhogal, B. et al. Modulation of dADAR-dependent RNA editing by the Drosophila fragile X mental retardation protein. Nat Neurosci 14, 1517–1524, doi:10.1038/nn.2950 (2011).

68 Shamay-Ramot, A. et al. Fmrp Interacts with Adar and Regulates RNA Editing, Synaptic Density and Locomotor Activity in Zebrafish. PLoS Genet 11, e1005702, doi:10.1371/journal.pgen.1005702 (2015).

69 Patzlaff, N. E., Nemec, K. M., Malone, S. G., Li, Y. & Zhao, X. Fragile X related protein 1 (FXR1P) regulates proliferation of adult neural stem cells. Human molecular genetics 26, 1340–1352, doi:10.1093/hmg/ddx034 (2017).

70 Penagarikano, O. et al. Absence of CNTNAP2 leads to epilepsy, neuronal migration abnormalities, and core autism-related deficits. Cell 147, 235–246, doi:10.1016/j.cell.2011.08.040 (2011).

71 O’Roak, B. J. & State, M. W. Autism genetics: strategies, challenges, and opportunities. Autism research: official journal of the International Society for Autism Research 1, 4–17, doi:10.1002/aur.3 (2008).

72 Kim, H. G. et al. Disruption of neurexin 1 associated with autism spectrum disorder. American journal of human genetics 82, 199–207, doi:10.1016/j.ajhg.2007.09.011 (2008).

73 Vaags, A. K. et al. Rare deletions at the neurexin 3 locus in autism spectrum disorder. American journal of human genetics 90, 133–141, doi:10.1016/j.ajhg.2011.11.025 (2012).

74 Lee, J.-A. et al. Cytoplasmic Rbfox1 regulates the expression of synaptic and autism-related genes. Neuron 89, 113–128 (2016).

75 Finucane, B. M. et al. in *GeneReviews(R)* (eds M. P. Adam et al.) (University of Washington, Seattle. GeneReviews is a registered trademark of the University of Washington, Seattle. All rights reserved., 1993).

76 Ahn, J. & Xiao, X. RASER: reads aligner for SNPs and editing sites of RNA. Bioinformatics (Oxford, England) 31, 3906–3913, doi:10.1093/bioinformatics/btv505 (2015).

77 Josse, J. & Husson, F. missMDA: A Package for Handling Missing Values in Multivariate Data Analysis. 2016 70, 31, doi:10.18637/jss.v070.i01 (2016).

78 Langfelder, P., Zhang, B. & Horvath, S. Defining clusters from a hierarchical cluster tree: the Dynamic Tree Cut package for R. Bioinformatics (Oxford, England) 24, 719–720, doi:10.1093/bioinformatics/btm563 (2008).

79 Wheeler, E. C., Van Nostrand, E. L. & Yeo, G. W. Advances and challenges in the detection of transcriptome-wide protein-RNA interactions. Wiley interdisciplinary reviews. RNA, doi:10.1002/wrna.1436 (2017).

80 Heinz, S. et al. Simple combinations of lineage-determining transcription factors prime cis-regulatory elements required for macrophage and B cell identities. Mol Cell 38, 576-589, doi:10.1016/j.molcel.2010.05.004 (2010).

81 Bailey, T. L., Johnson, J., Grant, C. E. & Noble, W. S. The MEME Suite. Nucleic acids research 43, W39–49, doi:10.1093/nar/gkv416 (2015).

82 Aken, B. L. et al. The Ensembl gene annotation system. Database: the journal of biological databases and curation 2016, doi:10.1093/database/baw093 (2016).

83 Nojima, T., Gomes, T., Carmo-Fonseca, M. & Proudfoot, N. J. Mammalian NET-seq analysis defines nascent RNA profiles and associated RNA processing genome-wide. Nat Protoc 11, 413–428, doi:10.1038/nprot.2016.012 (2016).

84 Sundararaman, B. et al. Resources for the Comprehensive Discovery of Functional RNA Elements. Mol Cell 61, 903–913, doi:10.1016/j.molcel.2016.02.012 (2016).

85 Bahn, J. H. et al. Genomic analysis of ADAR1 binding and its involvement in multiple RNA processing pathways. Nat Commun 6, 6355, doi:10.1038/ncomms7355 (2015).

